# Astrocyte 3D Culture and Bioprinting using Peptide Functionalized Hyaluronan Hydrogels

**DOI:** 10.1101/2022.10.13.510969

**Authors:** Isabelle Matthiesen, Michael Jury, Fatemeh Rasti Boroojeni, Saskia L. Ludwig, Muriel Holzreuter, Sebastian Buchmann, Andrea Träger, Robert Selegård, Thomas E. Winkler, Daniel Aili, Anna Herland

**Affiliations:** Division of Micro and Nanosystems, KTH Royal Institute of Technology, 10044 Stockholm, Sweden; CVRM Safety, Clinical Pharmacology and Safety Sciences, R&D, AstraZeneca, Gothenburg, Sweden; Laboratory of Molecular Materials, Division of Biophysics and Bioengineering, Department of Physics, Chemistry and Biology, Linköping University, 581 83 Linköping, Sweden; Institute of Microtechnology & Center of Pharmaceutical Engineering, Technische Universität Braunschweig, 38106 Braunschweig, Germany; AIMES, Center for Integrated Medical and Engineering Science, Department of Neuroscience, Karolinska Institute, 17165 Solna, Sweden; Division of Nanobiotechnology, Department of Protein Science, Science for Life Laboratory, KTH Royal Institute of Technology, Solna, 17165 Sweden

**Keywords:** astrocytes, 3D cell culture, bioprinting, hyaluronan, cRGD, IKVAV

## Abstract

The often-forgotten astrocytes play an important role in the central nervous system, contributing to the development of and maintenance of synapses, recycling of neurotransmitters, and the pathophysiology of various neurodegenerative diseases. Hydrogels can provide improved support and attachment for the culture of astrocytes in 3D models, which could further be used to advance clinical in vivo like tissue models of numerous diseases. For full applicability, these gels must be of scalable and defined origin and with stable attachment elements, such as peptides. In this study, the generation of a functional 3D astrocyte model is presented using a hyaluronan-based hydrogel system conjugated with the peptide sequences cyclic RGD (cRGD) and IKVAV, known promoters of cell attachment. Encapsulation of the neuroblastoma cell line SH-SY5Y and glioblastoma cell line U87 is successfully demonstrated over a 6-day culture period. The presence of the peptides cRGD and IKVAV does not change the cells’ viability. Human fetal primary astrocytes (FPA) are further tested for the 3D culture in these materials, similarly, showing that the peptides have no effect on the viability over a 6-day culture period. mRNA expression analysis reveals no biologically significant changes in the 3D cultures FPA or the U87 cells. Morphological analysis, on the other hand, revealed that FPA have a higher degree of interactions with the hyaluronan-based gels compared to the cell lines. This interaction is enhanced by peptide conjugation, in particular cRGD. Finally, we demonstrated that the peptide conjugated hydrogels could be used for bioprinting of FPA, opening up for defined neural astrocytic co-culture.

## Introduction

Of all the cells that make up the human brain, astrocytes are the most abundant cell type. The function of astrocytes was, for a long time, reduced to that of a glue-like scaffold helping to structure the brain tissue. However, recent studies provide evidence that astrocytes are involved in synapse formation and function, as well as metabolic activities to support neurons through glutamate clearance at the synaptic cleft [1–3]. Astrocytes have also been proven to be highly involved in several pathological conditions of the central nervous system (CNS); for example, they act protectively by producing and secreting the antioxidant glutathione during oxidative stress, which is a common trait in neurodegenerative diseases [4–6]. The size and the shape of astrocytes also vary during different disease conditions. When astrocytes were studied in an epileptic model, they extended thicker and longer protrusion than healthy cells [7]. During traumatic brain injury, where tissue lesions were formed, astrocytes were seen to extend long processes towards the injured area [8]. Such lesions have also been studied in 3D hydrogel models based on alginate gels, where glial marker GFAP expression changes were observed when cultured in meningeal fibroblast conditioned medium [9]. Alginate, collagen and other commonly used commercial scaffolds have been used to create various 3D models, however, lack available cellular attachment motifs. Collagen, an ECM component itself, does feature RGD motifs, but these are hidden within the structure[10]. Therefore, the design of hydrogel systems that are compatible with biofabrication techniques and simultaneously promote natural cell attachment is important for developing in vitro models of the CNS and other tissues. The short peptide motifs Arg-Gly-Asp (RGD) and Ile-Lys-Val-Ala-Val (IKVAV) are derived from ECM proteins such as laminins and have been proposed to improve cellular attachment in synthetic ECM models [11]. Integration of linear or cyclic RGD (cRGD) has previously been explored by us using a hyaluronan-based gel to better mimic the in vivo environment for the 3D culture of hepatoma cells and human induced pluripotent stem cell-derived hepatocytes in a perfused device [12]. Others have also used hyaluronan-containing RGD-functionalized hydrogels to investigate how the microenvironment contributes to cellular attachment and the survival of malignant brain tumor cells. The study showed that blocking the overexpressed hyaluronan receptor CD44 in the tumor cells reduces their adhesion to the ECM [13]. This motivates the development of 3D cellular models to enable detailed investigations of the mechanism of cellular attachment and spreading, which are important factors in the understanding of many disease models. The access to human-like ECM materials that are easy to use could also contribute to the development of more structurally advanced culture models, including 3D bioprinted architectures and microphysiological systems.

In this work, we present a bioprintable hyaluronan hydrogel system with the possibility of conjugating cell adhesion peptides (cRGD and IKVAV). The peptide functionalized hydrogels were assessed for the 3D culture of the widely used astroglia models SH-SY5Y cells and glioblastoma cell line U87. Furthermore, we evaluated the peptide conjugated hydrogel system for use in 3D cell culture of human fetal primary astrocytes (FPA) and as a bioink for 3D bioprinting of astrocyte seeded hydrogel structures.

## Experimental Section

### Synthesis of HA-BCN

HA (150 kDa) obtained from Lifecore Biomedical (Minnesota, USA) was modified with bicyclo[6.1.0]nonyne (BCN) as described in further detail in Selegård et al. [14]. In brief, N-[(1 R,8 S,9 S)-Bicyclo[6.1.0]non-4-yn-9-ylmethyloxycarbonyl]-1,8diamino-3,6-dioxaoctane (BCN-NH_2_) was dissolved in 5:1 (v/v) acetonitrile:MilliQ water prior to adding 1-Ethyl-3-[3-dimethylaminopropyl]carbodiimide hydrochloride and 1-hydroxybenzotriazole hydrate. The mixture was then added to HA previously dissolved in 2-(N-morpholino)ethanesulfonic acid buffer (100 mM, pH 7). The carbodiimide reaction was allowed to continue for 24 h on a shaker at room temperature. Subsequently, time dialysis (MWCO 12-14 kDa, Spectra/Por RC, Spectrum Laboratories Inc.) was performed in acetonitrile:MilliQ water (1:10 v/v) for 24 h and then for three days in MilliQ water. The final product, denoted as HA-BCN, was lyophilized and stored at -20°C. Based on 1H-NMR analysis, the HA-BCN had a derivatization degree of approximately 19%.

### Peptide synthesis

The peptide cRGD with the sequence c(RGDfK(N_3_) was synthesized as described previously [11]. The peptide IKVAV with the sequence K(N_3_)GGIKVAV-NH_2_ was synthesized on a Liberty Blue peptide synthesizer (CEM, Matthews, USA) using standard fluorenylmethoxycarbonyl protecting group (Fmoc) chemistry. ProTide Rink Amide (0.19 mmol/g) was used as solid support, and the peptide was synthesized in a 100 μmol scale. Each coupling was performed under microwave conditions at 90 °C for 2 min using a fivefold excess of amino acid and N,N-diisopropylcarbodiimid as coupling reagent and tenfold excess of Oxyma Pure as base. Fmoc deprotection was also performed under microwave conditions at 90 °C for 1 min using 20% piperidine in dimethylformamide (DMF) (v/v). The peptide was, after final Fmoc deprotection, cleaved from its solid support by treatment with trifluoroacetic acid (TFA):H_2_O:trisioproylsilane (95:2.5:2.5, v/v/v) for 3 hours before being concentrated and precipitated using ice-cold diethyl ether. The crude peptide was purified with a ReproSil gold C-18 column attached to a semi-preparative HPLC system (UltiMate 3000, Dionex, Sunnyvale, USA) using an aqueous gradient of acetonitrile with 0.1% TFA. Identity of the peptide was confirmed using a MALDI-ToF mass spectrometer (UltrafleXtreme, Bruker, Billerica, USA) running in positive ion mode with alpha-cyano-4-hydoxycinnamic acid as matrix (Figure S2). Peptide purity was confirmed using HPLC (Figure S3).

### Hydrogel formation

HA-BCN and 8-arm Az-terminated polyglycol ethylene ((PEG-Az)_8_) was suspended in cell media (including, where relevant, cells) unless stated otherwise to give a concentration of 1% (w/v). 10 mM peptide solution was added to the HA-BCN as 10% of total hydrogel volume (i.e., 1 mM final concentration) and incubated for one hour at 37°C to allow for HA functionalization via strain-promoted alkyne-azide cycloadditions (SPAAC). The same volume of PBS was added instead of the peptide in the no peptide condition. HA-BCN + peptide and (PEG-Az)_8_ were combined at a ratio of 3:1, quickly but thoroughly mixed, and the gels formed as required. The plate was sealed, and incubation was performed for 1 hour at 37°C to cross-link the gels, after which pre-warmed cell media was added to the wells.

### Rheology

Oscillatory rheology was performed using a Discovery HR-2 rheometer (TA Instruments, USA). Hydrogels were produced with a total volume of 30 μl as described above. Instead of cell media, PBS was added to completely cover the hydrogels and incubation continued for a further 24 hours at 37°C. The hydrogels were analyzed using strain sweeps (1 Hz, 0.1 to 10% strain) and frequency sweeps (1% strain, 0.1 to 10 Hz) using an 8 mm parallel geometry. Gelation kinetics were performed using 50 μl volumes and a 20 mm 1° geometry at 1 Hz and 1% strain. Hydrogel components were mixed immediately prior to measurement and added to the platter precooled to 4°C. With the geometry in place, the platter was rapidly warmed to 37°C and measurements began. It is estimated that 10 -20 seconds passed between mixing components and measurements starting.

### Scanning Electron Microscopy (SEM)

Hydrogels were produced as described, adding phosphate buffer (PB, 10 mM, 7.4 pH) instead of cell media to completely cover the hydrogels after cross-linking. Incubation at 37°C was continued for a further 24 hours. The resulting hydrogels were carefully placed onto carbon disc sample holders, rapidly frozen with liquid nitrogen and lyophilized. The dried samples were carefully sliced to expose the internal structure and then sputter-coated with platinum for 10 seconds. SEM analysis was conducted with a LEO 1550 Gemini (Zeiss, Germany) operating as a voltage of 3 kV.

### SH-SY5Y cell culture

SH-SY5Y cells were obtained from ATCC, USA (CRL-2266) and were thawed and cultured as per the supplier’s instructions in DMEM/F12 (1:1) cell media (Biowest, USA) supplemented with 10% FBS (Biowest, USA), 1% PenStrep (Biowest, USA) and 1% non-essential amino acids (Biowest, USA), hereby denoted as SH-SY5Y maintenance media. Cells were cultured in T75 flask and split 1:4 when confluence exceeded 80%. Passage number was never allowed to exceed 20 above the passage number at which the cells were supplied.

### Viability of SH-SY5Y cells

Hydrogel components were prepared using SH-SY5Y maintenance media as the carrier solution. Confluent SH-SY5Y cultures were trypsinized (Trypsin 0.25% in PBS, Biowest, USA) for 2 minutes at 37°C with gentle agitation to ensure a homogenous single-cell suspension and counted with trypan blue and a Bürker chamber. Counted cells were pelletized at 120g for 6 minutes and then suspended in the (PEG-Az)_8_ component. =Hydrogels were formed as described above, with a cell count of 10^5^ per 50 μl hydrogel volume. Cells were cultured for up to 6 days at 37°C and 5% CO2. At time points of 1, 3 and 6 days, the media was removed, and wells gently washed with PBS. AlamarBlue™ Cell Viability Reagent (AB) (Invitrogen, USA) was used as the viability assay, with 10% added to cell media and placed in each well. Incubation followed for two hours at 37°C and 5% CO2, after which the AB solution was moved to a new 96 well plate for measurement of absorbance at 570 nm, and 600 nm (±5nm) using a TECAN infinite M1000 Pro (Tecan, Switzerland). The absorbance measurements were calculated into percentage reduction as an indicator of cell viability using the calculation set out in the official protocol.

### FPA and U87 cell culture

All reagents were purchased from Thermo Fisher Scientific, MA, USA unless stated otherwise. Human primary fetal astrocytes here, referred to as FPA (ScienCell, USA), were maintained as per the supplier’s instructions. In brief, the cells were cultured in tissue culture-treated T75 flasks coated with 1x Attachment Factor protein in ScienCell astrocyte media (AM) with the supplements of 2% FBS, 1% astrocyte growth supplement (AGS) and 1% PenStrep (all ScienCell). When confluent, cells were passaged at a ratio of 1:4, using TrypLE incubation for 4 min. Defined Trypsin Inhibitor (DTI) was used for TrypLE inhibition; the cells were spun down to a pellet at 200g, after which they were resuspended in the supplemented AM. The U87 glioblastoma cells, previously published by Pontén and Macintyre^[15]^ and obtained from Uppsala University, were maintained in tissue culture-treated T75 flasks in high glucose DMEM supplemented with 10% FBS and 1 % PenStrep. When confluent, cells were passaged at a ratio of 1:4 by a 4 min TrypLE incubation. The cells were then spun down to a pellet at 200 g, after which they were resuspended in the supplemented DMEM described above.

### Encapsulation of FPA and U87 cells

Two days before encapsulation of the cells into 3D hydrogels, both the FPA and the U87 were gradually adjusted to a serum-free media. Serum-free media made from AM supplemented with 1% G5 supplement, 1% AGS and 1% PenStrep was mixed 1:1 with respective maintenance media of each cell type, upon which the media was given to the cells. One day before encapsulation, the media of the FPA and the U87 was changed fully to the serum-free media. 1.6 million cells/ml of FPA and U87 were encapsulated in 1% HA:PEG hydrogels containing 1 mM of either peptide and carefully pipetted into half-area 96 well plates in gels of 50 μl each. The cells were cultured for six days and compared to 2D cultures on standard tissue culture plates. The respective peptides’ influence was studied with AB viability measurement after 1, 3 and 6 days of culture in the hydrogels. After six days of subculture, the cells were lysed, and mRNA expression of the intermediate filament protein Vimentin (VIM), the focal adhesion protein Paxillin (PAX), integrin subunit beta 1 (ITGB1), the focal adhesion kinase Protein Tyrosine Kinase 2 (PTK2) and the hyaluronan receptor CD44 were investigated for both cell types with qPCR. Glial fibrillary acidic protein (GFAP) and calcium-binding protein B (S100B) were additionally assayed for the FPA. The samples were studied as technical duplicates and contained the gene of interest (GOI) and the chosen housekeeping gene (HKG) GAPDH. Expression of the GOI is presented as *ΔΔCt* normalized to the reference sample (RS), in this case, 2D-cell cultures, according to the following calculation:

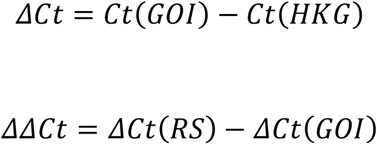

Origin Pro was used to calculate p-values using Linear Mixed Models (LMM). Any samples measured to a Ct higher than 35 and technical duplicates with a variation higher than one were excluded.

Both cell types were stained for CD44, F-actin, and nuclei to visualize the cellular morphology and imaged with a confocal microscope. The confocal images were composed by maximum projection, after which brightness and contrast were adjusted to accentuate best the structures illustrating the cellular morphology. The image analysis was carried out in Fiji ^[16]^. To quantify the actin filament spreading and branchpoints the FIJI macro TWOMBLI was used with the following parameters: contrast saturation 0.35, line width 15, curvature window 50, minimum branch length 10, and minimum gap diameter 0 ^[17]^.

### 3D bioprinting

All 3D bioprinting was performed using a syringe extrusion printhead on a Cellink BIO X 3D bioprinter (BICO, USA). FPA were cultured until confluent and then detached using TrypLE express. The cells were suspended in the (Peg-Az)_8_ component at a concentration of 200 000 per 100 μl of a final hydrogel. Hydrogel components were prepared and combined as described above, with the exception that Cy5-azide was added at a concentration of approximately 40 μM to facilitate imaging of bioprinted structures. Hydrogel grid structures were printed directly on sterile tissue-culture treated plates, sealed and incubated at 37°C and 5% CO_2_ for one hour for gelation to occur. After one-hour, pre-warmed co-culture media was added, and the plates were cultured for four days, after which the grid structures were fixated with 0.5% paraformaldehyde in preparation for staining.

## Results and Discussion

### Hydrogel formation

Hyaluronan (HA) based hydrogels were obtained using copper-free klick-chemistry as described previously, ^[12,18]^ using bicyclo[6.1.0]nonyne (BCN) modified HA (≈100 kDa) and an 8-arm PEG with terminal azide (Az) groups ((PEG-Az)_8_) and as a cross-linker (Figure 1d). Rheological characterization of the resulting hydrogels using strain and frequency sweeps (Figure 1a,b) shows that the hydrogels were indeed cross-linked, also after conjugation of adhesion peptides, as indicated by a storage modulus (G’) higher than loss modulus (G”) for all conditions in the linear-viscoelastic region and absence of cross-over points. The G’ of the non-peptide functionalized HA hydrogels was about 175 Pa at 1 % strain and 1 Hz. Electron micrographs (Figure 1b) show a microporous structure, which can facilitate cell migration in the hydrogels. The conjugation of cell adhesion peptides cRGD and IKVAV to the HA backbone resulted in a decrease in the stiffness of the hydrogels to about 72 and 118 Pa, respectively (Figure 1a, b). Since peptide conjugation reduces the number of BCN groups available for cross-linking, a slight reduction in stiffness is expected. Both peptides were conjugated to a final overall concentration of 1 mM within the hydrogels. The difference in stiffness after conjugation of the two peptides could indicate a slightly more efficient conjugation of cRGD compared to IKVAV; however, differences in chemical properties of the two peptides could also contribute to these effects. Interestingly, the decrease here is in contrast to the insignificant change in stiffness we observed in our prior study with conjugation of azide-functionalized full-length recombinant laminin, where the large protein likely contributed to the cross-linking of the hydrogels.[1] Regardless, our HA:PEG gel stiffness is within the typical range for neural tissue [19].

**Figure 1.**
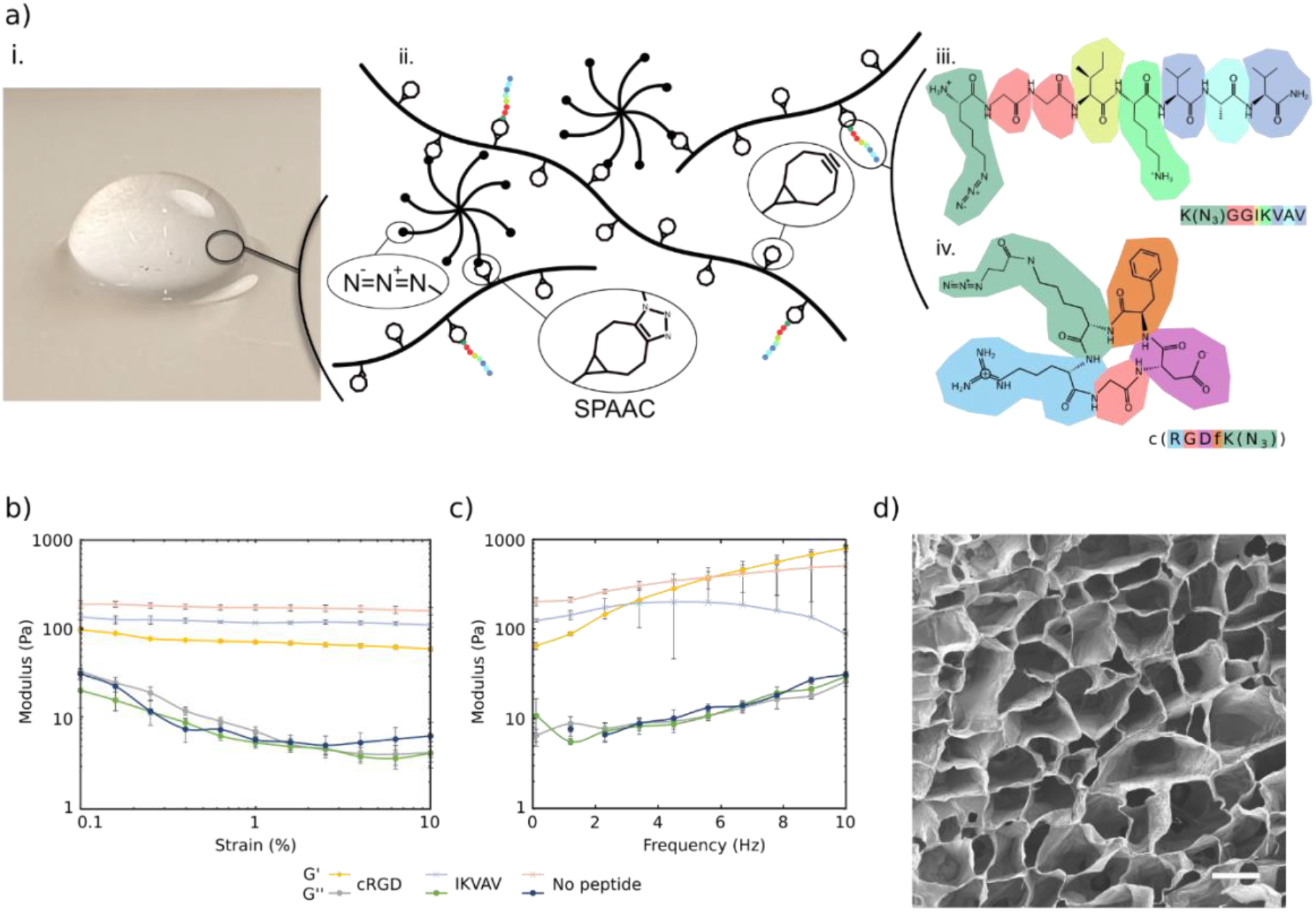
a) i. Photographic image of hyaluronan-based hydrogel. ii. schematic illustration of the HA:PEG: peptide system with the detailed amino acid structures of the peptides iii. IKVAV and iv. cRGD. b) Strain sweeps from 0.1 to 10 % and at 1 Hz of hydrogels, and c) frequency sweeps from 0.1 to 10 Hz and at 1 % strain of hydrogels with 1% (w/v) HA-BCN:(PEG-Az)_8_. Error bars represent standard deviation (n=3). d) SEM image of lyophilized 1% (w/v) HA-BCN:(PEG-Az)_8_ hydrogel. Scale bar is 50 μm.

### SH-SY5Y viability

For initial evaluation of hydrogel biocompatibility, we employed the widely-used SH-SY5Y neuroblastoma cells and assessed their viability using Alamar blue (AB) at time points of 1, 3 and 6 days (Figure 2a). The percentage of reduction of AB was used as an indicator for cell viability since it indicates viable metabolizing cell populations. At the day 1 time point, there was a significant increase in metabolic activity between the cRGD and no peptide condition; however, there was no significance between the IKVAV and the no peptide condition for the same time point. After six days in culture, all three conditions showed a significant decrease in metabolic activity. The 3D cell cultures were then fixated and stained for F-actin (Figure 2b) at time points 1 and 6. The representative images show that the cells in the cRGD functionalized hydrogels were sparser but were equally less prone to form large spheroids or cell clusters. Such cell clustering is evident in the IKVAV-functionalized hydrogel and even more so in the no peptide condition. Clustering of cells is a known trait for the SH-SY5Y cell line and does not necessarily indicate an unviable cell population ^[20]^. Further, while a decrease in metabolic activity was observed over time, the micrographs show an increase in cell density and cell proliferation and less dense clusters of cells. The hydrogels thus provide adequate and similar support for cell proliferation irrespectively of peptide functionalization.

**Figure 2.**
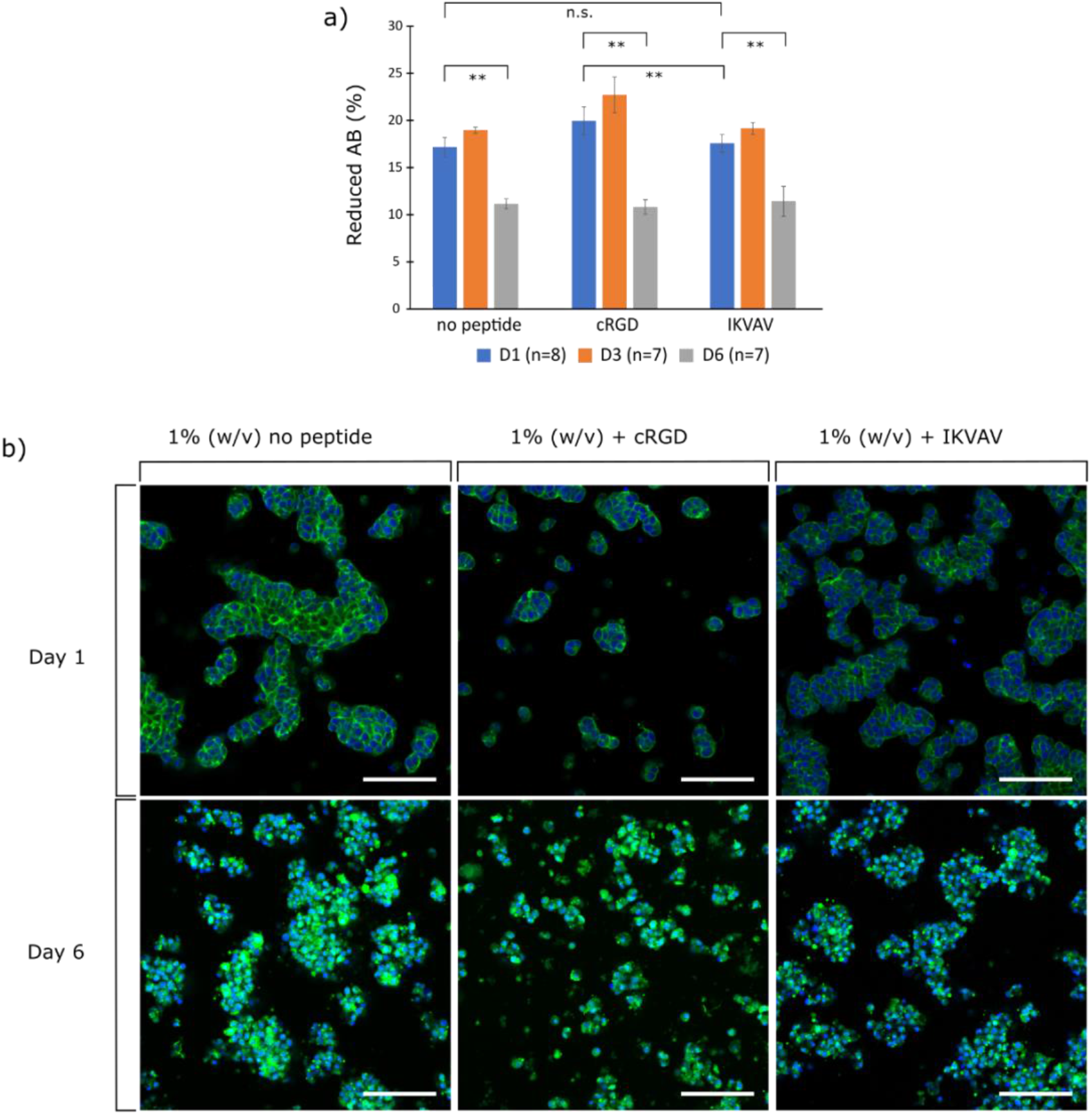
a) Cell viability of SH-SY5Y cell cultures using AlamarBlue™ Cell Viability Reagent (AB). Values correspond to the percentage of AB reduced by viable cells on day 1 and 6. Error bars are standard deviation and significance is calculated using ANOVA with Tukey HSD (n.s. = no significance/>0.05), ** = <0.01). b) Micrographs of 3D cell cultures of SH-SY5Y cells in 1% (w/v) HA-BCN:(Peg-Az)_8_. Green is phalloidin staining for F-actin, blue is Hoechst staining for nucleus. Scale bar = 100 μm.

### Viability of encapsulated FPA and U87 cells

To assess the possibility to support astrocytic cells primary human astrocytes were encapsulated in 3D hydrogels. Viability was studied using the AB viability assay. After 1 day of subculture, FPA show comparable viability in all gel conditions, Figure 3a. The addition of cRGD or IKVAV did not affect the viability at this stage.

**Figure 3.**
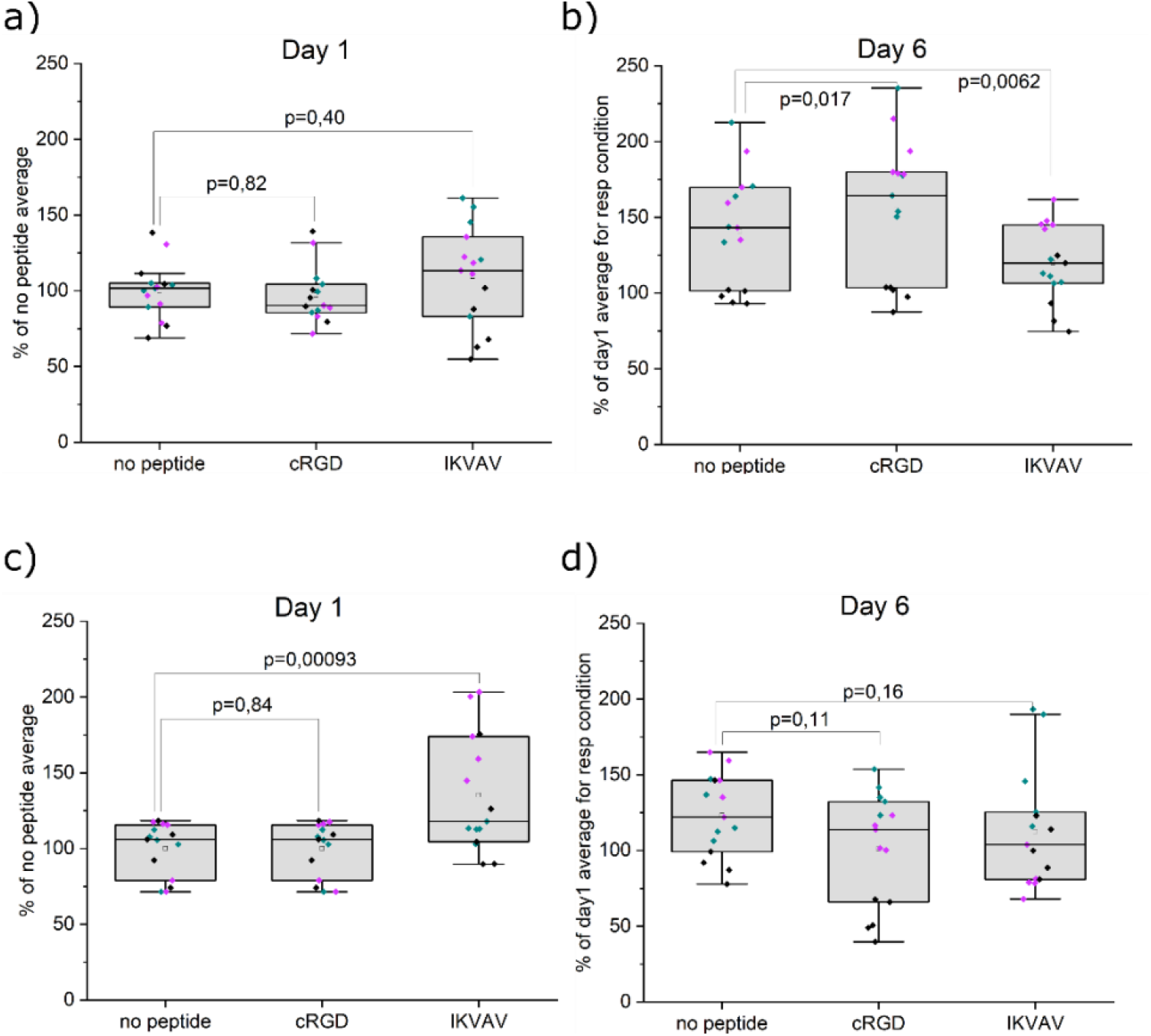
AB viability assay of encapsulated cells. a) Viability of FPA after 1 day in respective hydrogel conditions. b) Change in viability of FPA after 6 days in respective hydrogels relative to day 1. c) Viability of U87 after 1 day in respective hydrogel conditions. d) Change in viability of U87 after 6 days in respective hydrogels relative to day 1. Data were collected from three individual experiments (indicated with different colors. N=5, where N represents a hydrogel replicate. Origin Pro was used for derivation of p-values with the LMM method, all data included. Data are presented both as a box indicating the 25th–75th percentile, including a median line and ±1.5 IQR whiskers, with the addition of individual data points.

Over the course of 6 days (Figure 3b), all conditions increase in viability compared to the corresponding hydrogel condition at 1 day of subculture. Moreover, a significant increase in viability compared to the no-peptide HA is observed when cRGD is added to the hydrogel. Surprisingly, the addition of IKVAV leads to a less increase in AB reduction day 6 versus day 1 compared to the condition where no peptide was added. Regardless, these results indicate that the hydrogels are suitable for maintaining FPA; however, the peptide functionalization has limited effect on promoting proliferation. These results align with our previous observations for culture and differentiation of neural cells [1]. In addition to culture of primary astrocytes, we explored the often-used astrocytic model glioblastoma cell line U87. AB assay after 1 day of subculture shows viable cells in all gel conditions. The addition of IKVAV to the hydrogels resulted in a significant increase in viability here, while the addition of cRGD had no effect. When the AB assay was performed after 6 days of subculture, the results showed that the viability remained stable for all conditions, without significant effects from addition of cRGD or IKVAV. In broad terms, the patterns of U87 viability are not dissimilar from the FPA ones, but they are a clear contrast to the SH-SY5Y. With the neuroblastoma cells, cRGD rather than IKVAV improved short-term viability, and viability decreased significantly by day 6 for all conditions. This suggests that the different cell lineages rely on distinct dominant mechanisms for interaction with the hydrogel.

### *mRNA expression* analysis *of encapsulated FPA and U87*

After 6 days of subculture in 3D hydrogels, the cells were lysed, and mRNA expression levels were measured and normalized to the cells cultured in conventional 2D tissue culture plates. (Figure 4, Figure 5). Seven attachment-or astrocytic-related markers were tested for FPA, out of which five markers were also tested for the U87 cells as they do not express all astrocytic-related markers. To begin with, the integrin ITGB1 was assessed and was detected in both cell types, with no difference in expression for any of the hydrogel conditions or cell types (Figure 4 a, Figure 5 a) with p-values presented in Table S1 a) and b). Similarly to ITGB1, the intermediate filament protein Vimentin (VIM), the focal adhesion protein Paxillin (PAX), and the focal adhesion kinase Protein Tyrosine Kinase 2 (PTK2) showed no change in expression between non-functionalized hydrogels and hydrogels functionalized with cRGD or IKVAV (Figure 4 and 5 b-d), see Table S1 for p-values. Importantly, no differences in hyaluronan-binding CD44 expression were seen between the different hydrogel conditions (Figure 4 e, Figure 5 e). Attempts to block the dimerization of CD44 have previously been observed to be an efficient way of controlling cellular attachment in tumors, suggesting that such studies can be interesting for understanding astrocyte adhesion to the HA:PEG hydrogels used here^[21,22]^. We investigated blocking CD44-gel interactions with either antibodies or a small molecule-compound; however, surprisingly, the data did not support a major change in hydrogel interactions (data not shown). However, a larger panel of CD44 blocking methods would be needed to fully exclude this potential cell-gel interaction. Furthermore, two astrocytic markers were assayed for FPA, namely the calcium-binding protein B (S100B) and the glial fibrillary acidic protein (GFAP). No difference in mRNA expression levels were seen upon functionalization of the hydrogels with either IKVAV or cRGD (Figure 4 f-g), indicating that astrocyte phenotype was not influenced by additional integrin-mediated interactions between FPA and the hydrogels. Broadly speaking, expression of all studied genes thus remains conserved across the 2D and 3D culture systems studied here and suggests that we have a stable *in vitro* phenotype in both conditons.

**Figure 4.**
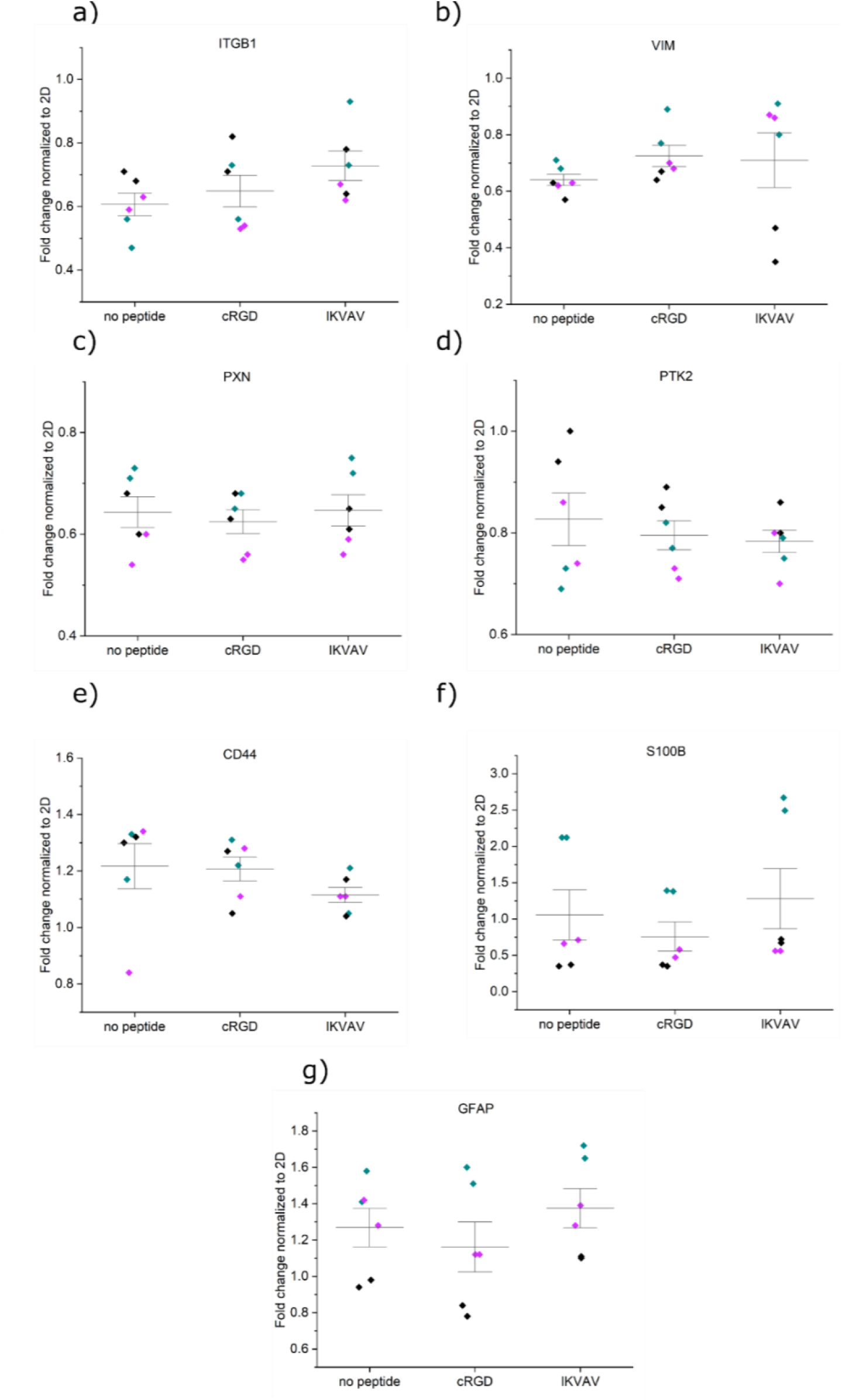
mRNA expression analysis of encapsulated FPA in different hydrogel conditions a) ITGB1, b) VIM, c) PXN, d) PTK2, e) CD44, f) S100B, and g) GFAP. All samples contain the housekeeping gene GAPDH. Origin Pro was used for the derivation of p-values with the LMM method. Data were collected as duplicates from three independent experiments, indicated with different colors.

**Figure 5.**
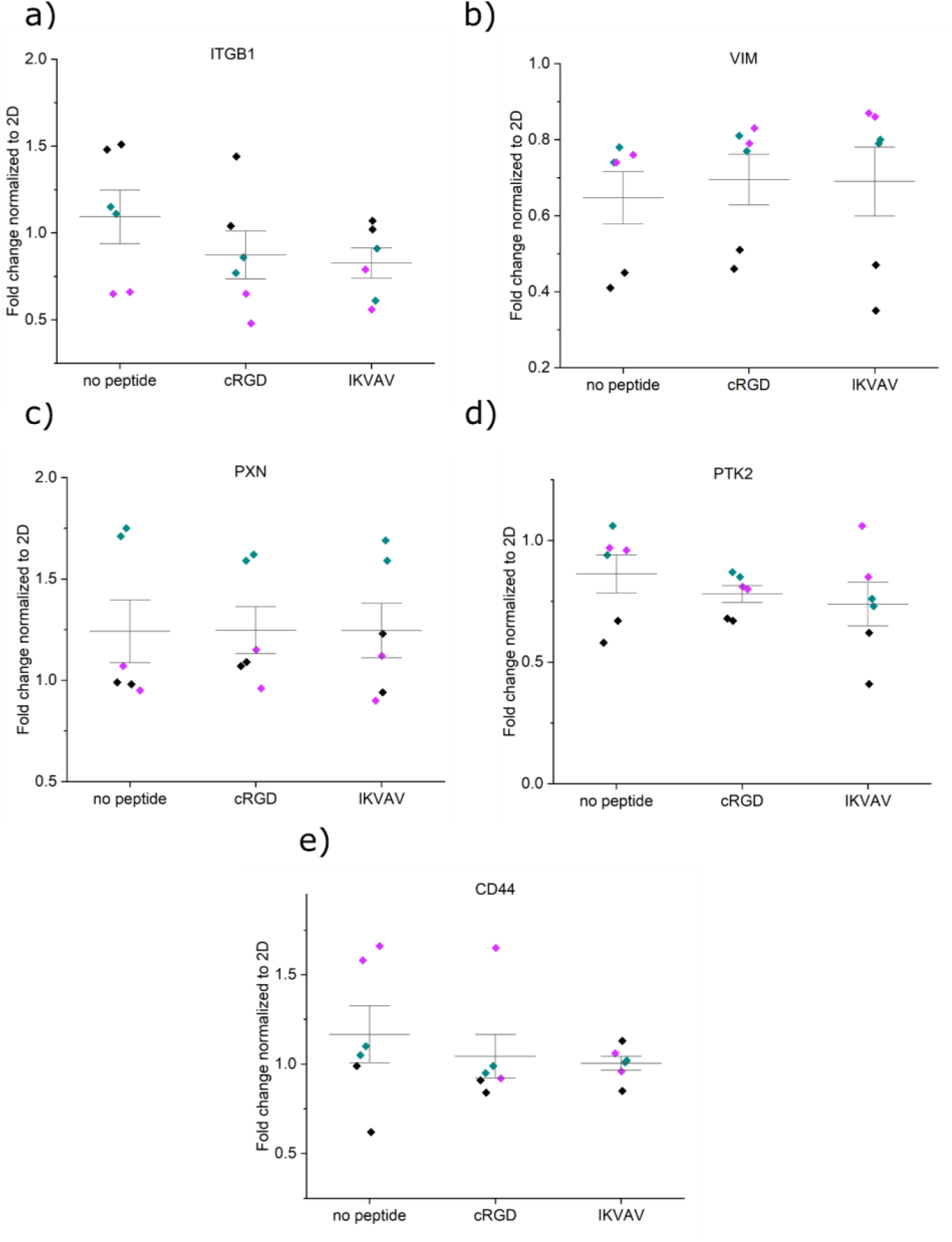
mRNA expression analysis of encapsulated U87 in different hydrogel conditions a) ITGB1, b) VIM, c) PXN, d) PTK2, and e) CD44. All samples contain the housekeeping gene GAPDH. Origin Pro was used for the derivation of p-values with the LMM method. Data were collected as duplicates from three independent experiments, indicated with different colors.

### Morphology of encapsulated FPA and U87 cells

The interactions between the cells and the hydrogels were further studied by confocal microscopy and immunostaining of the hyaluronic acid-binding CD44 receptor and F-actin staining for visualization of cytoskeletal organization (Figure 6). 3D encapsulated FPA cultured in hydrogels without peptides appeared to some extent to grow as single cells with a slightly condensed soma, while other cells spread out their protrusions and connected with neighboring cells and attached to the hydrogel (Figure 6a). CD44 is partially located as a surrounding halo of the nuclei and expressed along with the soma and cellular protrusions. Similarly, F-actin staining revealed a rounded shape of a subpopulation of the cells, while other cells appeared more stellate and interacting with the hydrogel matrix and other cells. Cell nuclei appeared oval and healthy under all conditions. A minority of the cells showed condensed nuclei with a bright and smaller appearance. The conjugation of cRGD or IKVAV to the hydrogel resulted in a slightly different morphology where cells extended their protrusions and spread out to a greater extent, seen as a larger surface area of the individual cells and more extensive cell-cell contacts. The localization of CD44 was similar in both the cRGD and IKVAV conditions and was seen both in the soma and along the cellular extensions. F-actin staining show in detail how the cells spread out and connected to other cells nearby.

**Figure 6.**
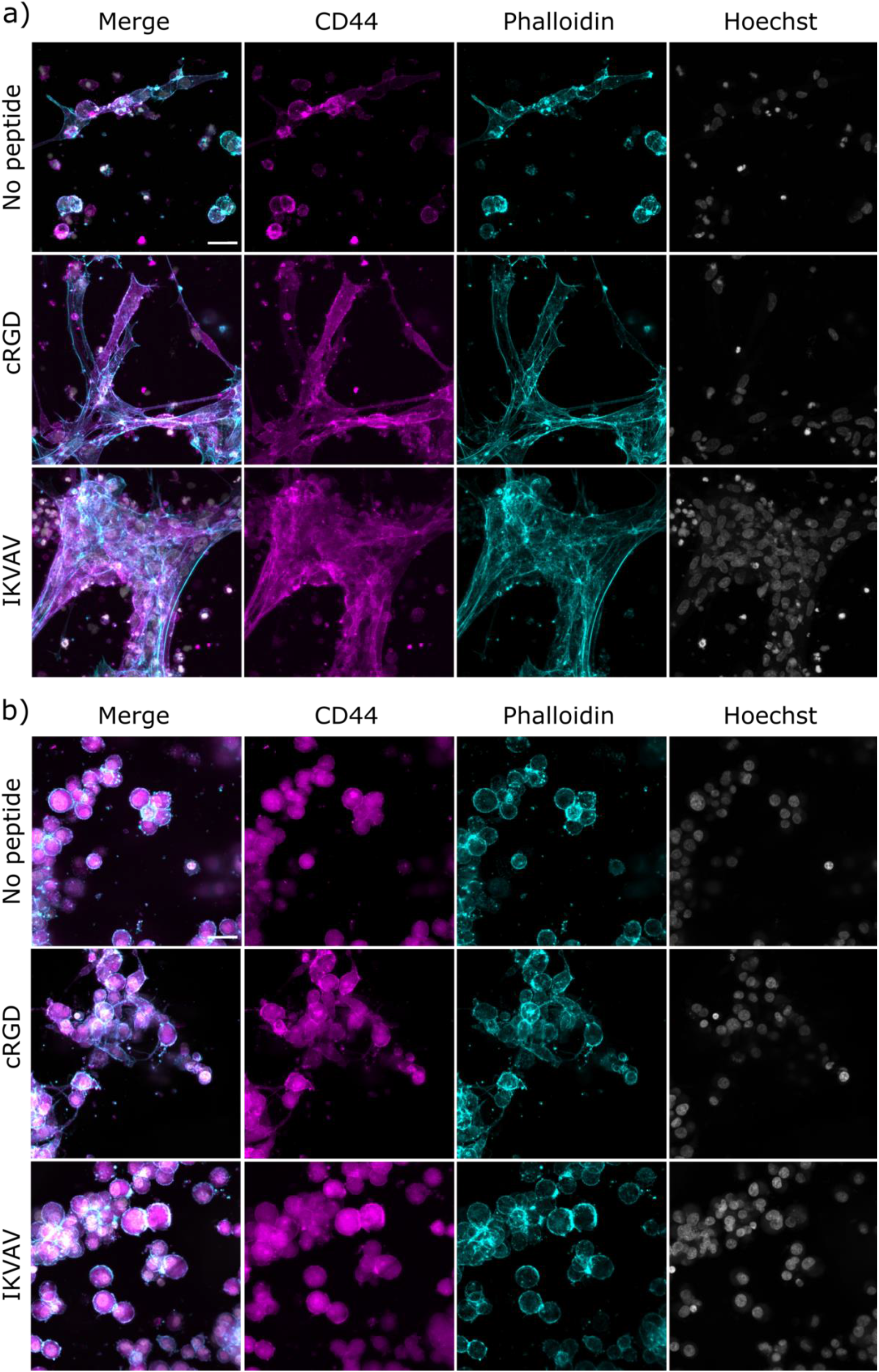
Confocal images of encapsulated FPA (a) and U87 (b) in respective hydrogels. FPA and U87 are stained with hyaluronic acid receptor CD44 (magenta), cytoskeletal F-actin marker Phalloidin (cyan), and nuclear stain Hoechst (gray). Scale bar: 30μm.

In contrast to the extensive spreading of FPA in the hydrogels, U87 cells typically appeared clustered and rounded; overall very similar to SH-SY5Y. All hydrogel conditions resulted in cells positive for CD44. The localization was limited to an area surrounding the cell nuclei (Figure 6b). The only exception was seen for the cRGD condition, where small extended intercellular connections were observed between certain cells. Similarly, F-actin staining visualized the partially condensed soma and little or no visible protrusions or cell-cell connections, apart from in cRGD functionalized hydrogels where some connections between cells were observed. Nuclei for all hydrogel conditions appeared healthy, although with a rounder shape than what was observed for FPA cultured under identical conditions.

We further employed quantitative image analysis of the F-actin networks (in terms of area per cell, and branch points per cell) to expand on our qualitative analysis above. These data fully support the notion that the FPA spread and branch out significantly more than the U87 across all gel conditions (Figure 7a-b). IKVAV, interestingly, shows no significant impact on these parameters compared to controls for either cell type. The contrast with our qualitative image observations may be attributed to the observed FPA networks in IKVAV-conjugated hydrogels being accompanied on also by larger cell numbers than the cRGD conditions. The data (Figure 7c-d) do in fact support a higher variability of morphology, which is also evident in Figure 6a and Figure S4 and which correlates with the distinct pattern in cell viability that IKVAV promoted in these cells (Figure 3a-b). Specifically, a higher variability of day 1 FPA viability, combined with lowered increase in viability toward day 6, would potentially support robust network formation in some cells. Still a large majority of cells remain condensed. With U87 (Figure 7e-f), a comparison with IKVAV viability data indicates that adhesion motifs supporting survival and proliferation do not necessarily translate to increased cell spreading and branching. The most notable finding from the analysis, however, is that in agreement with qualitative assessment, the addition of cRGD successfully and significantly promotes both the spreading and branching of either cell type.

**Figure 7.**
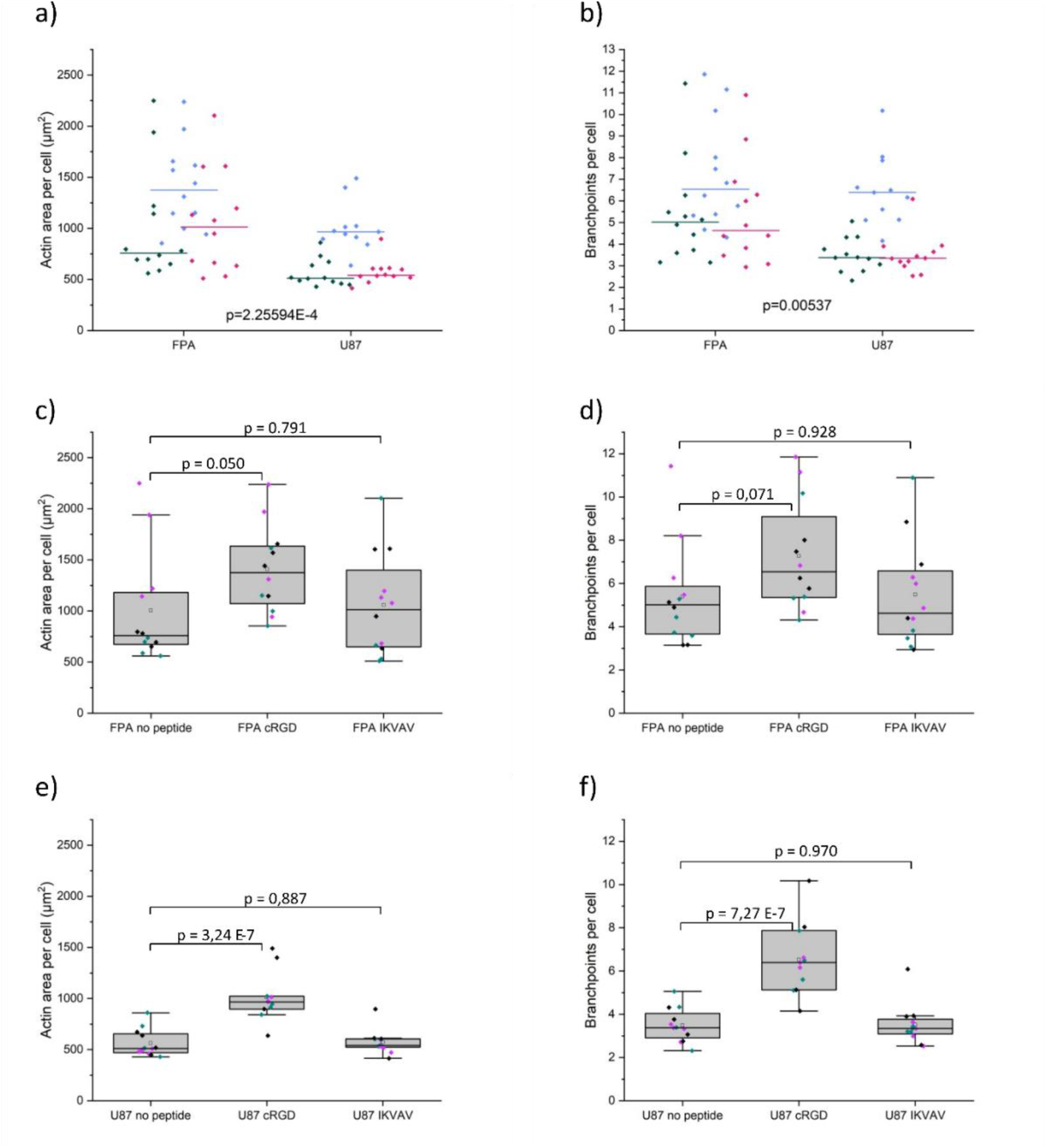
Quantification of actin filament spreading and branchpoints in FPA and U87 in respective hydrogels. a) Actin area per cell across all gel conditions. b) Branchpoints per cell across all gel conditions. In a-b) green is the condition no peptide, blue is cRGD and magenta is IKVAV c) Actin area per FPA cell. d) Branchpoints per FPA cell. e) Actin area per U87 cell. f) Branchpoints per U87 cell. Data were collected from three individual experiments N=5, where N represents a hydrogel replicate. In c-f) the different colors represent the different individual experiments. Origin Pro was used for derivation of p-values with the LMM method, all data included. Data are presented both as a box indicating the 25th–75th percentile, including a median line and ±1.5 IQR whiskers, with the addition of individual data points.

### 3D bioprinting

To finally investigate the possibilities of using the astrocyte-laden hydrogels as a bioink for 3D bioprinting, we printed hydrogel structures in a grid pattern, followed by culture for four days before investigating cell number and morphology (Figure 8). Despite being rather soft, our HA:PEG bioinks could be printed with good resolution. No apparent differences in cell density could be seen for the different hydrogel compositions. Cells printed using hydrogels without peptides grew as single rounded cells with a slightly condensed soma. Compared to gel casting, the even more condensed morphology is likely a result from added stress from the bioprinting process. However, presence of cell adhesion motifs resulted in cells with defined protrusions and multiple connections between neighboring cells in peptide-functionalized bioinks. With IKVAV, the difference compared to control appears more distinct here than in cast gels, potentially reflecting an increased relevance of the presence of any adhesion motif for pre-stressed cells. Branching and spreading is overall superior, however, with cRGD, in line with earlier observations.

**Figure 8.**
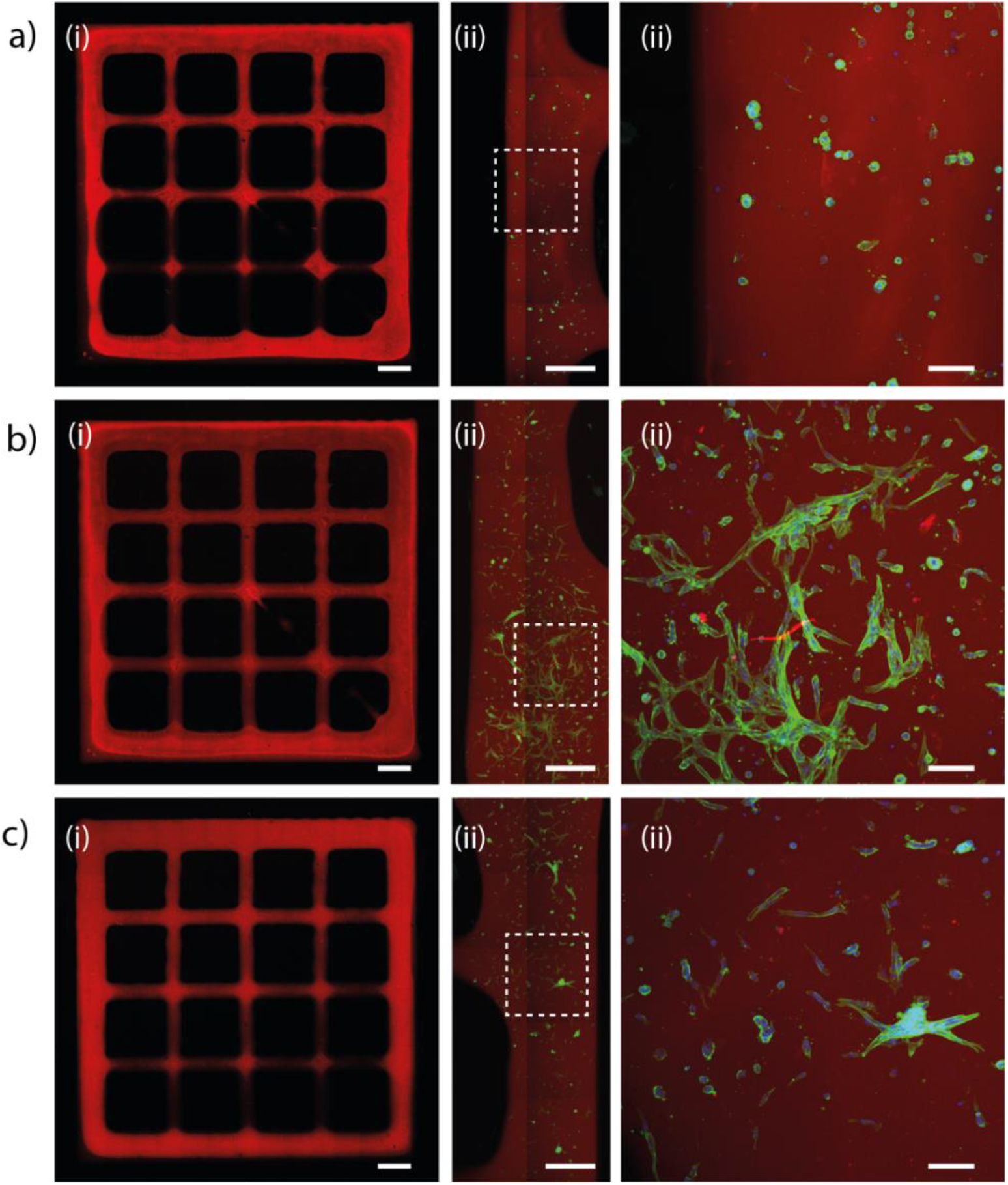
Bioprinted FPA at Day 4. The cells were encapsulated in HA-BCN:(PEG-Az)_8_ ± cRGD/IKVAV. a) HA-BCN:(Peg-Az)_8_. b) HA-BCN:(PEG-Az)_8_ functionalized with cRGD. c) HA-BCN:(PEG-Az)_8_ functionalized with IKVAV. The HA backbone was labelled with Cy5-Az (red), and FPA were stained green with phalloidin for F-actin. i) Full grid structure, scale bars are 1 mm. ii) Zoom in on part of the structure, scale bars are 0.5 mm. iii) Zoomed in view as indicated with white rectangles, scale bars are 100 μm. The images are representative of a larger set of imaged grid structures (N = 4 – 8).

## Conclusions

This study presents evaluation of 3D culture of cell models SH-SY5Y and U87 glioma cells and human FPA in a modular hyaluronan-based hydrogel system. The cells were cultured over a 6-day period. All three cell models express the hyaluronan binding receptor CD44, but display distinct morphologies. FPA are seen to spread out their processes and connect to neighboring cells in a network forming pattern, particularly in hydrogels functionalized with the fibronectin-derived cRGD-peptides. In contrast, U87 and SH-SY5Y cells mainly form clusters of cells showing little or no large-scale network formation under any of the conditions, though cRGD does show some benefits here as well. mRNA expression analysis of attachment and astrocytic markers display no difference between cells cultured in 2D or in hydrogels functionalized with IKVAV or cRGD. Bioprinting of FPA results in hydrogel dependent morphologies. Bioinks comprising hydrogels without peptide-cell adhesions motifs appeared rounded whereas cells in hydrogels modified with IKVAV and especially cRGD showed branched morphologies with extensive cell-cell contacts 4 days post printing. Successful bioprinting of FPAs open for development of more elaborate neural astrocytic co-culture models using this hyaluronan-based hydrogel system with defined cell adhesion motifs.

## Acknowledgments

I.M and M.J contributed equally to this work. The financial support from the Swedish Foundation for Strategic Research (SFF) (FFL15-0026), and the Knut and Alice Wallenberg Foundation (KAW 2016.0231, 2021.0186) is gratefully acknowledged.

## SUPPORTING INFORMATION

**Figure S1:**
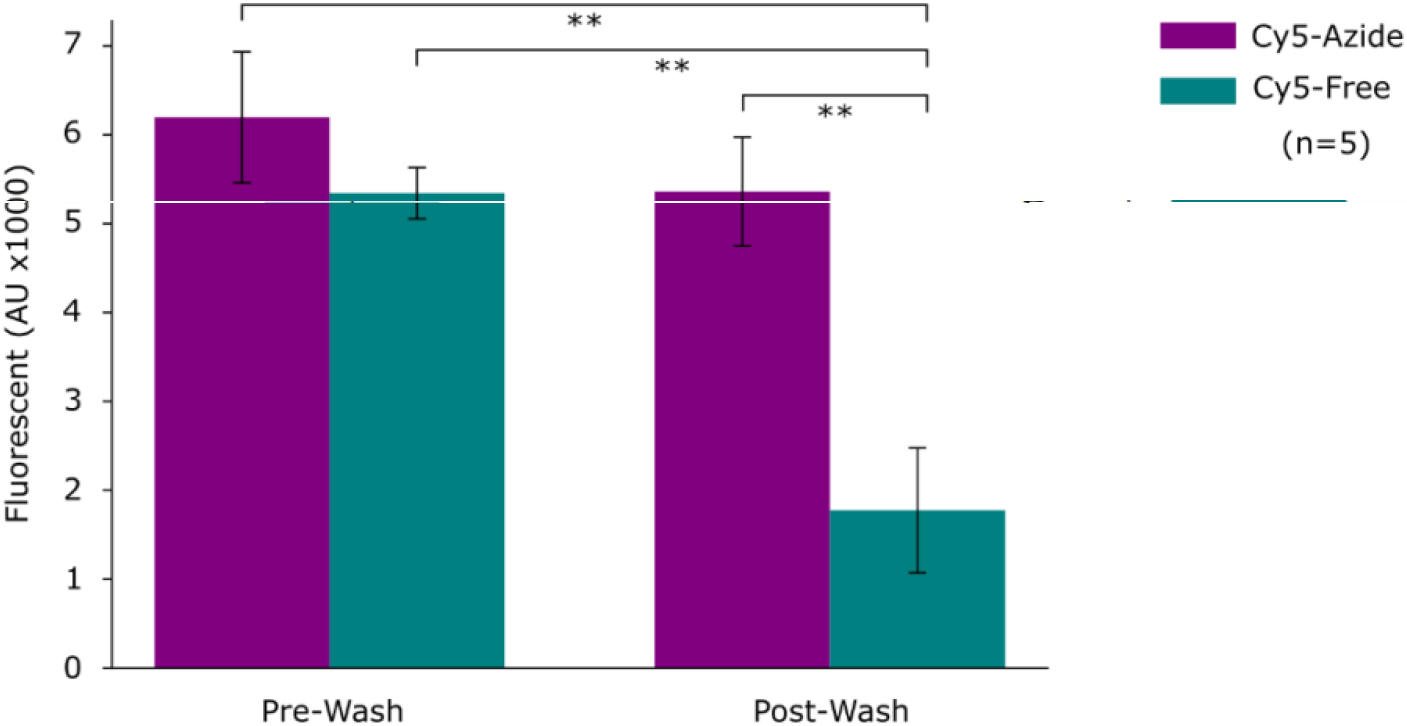
HA-BCN:(Peg-Az)8 hydrogels containing Cy5-azide and Cy5-free acid measured with a plate reader before and after washing. Significant release of Cy5-free acid (** = <0.01) and no significant (>0.05) release of Cy5-azide.

**Figure S2:**
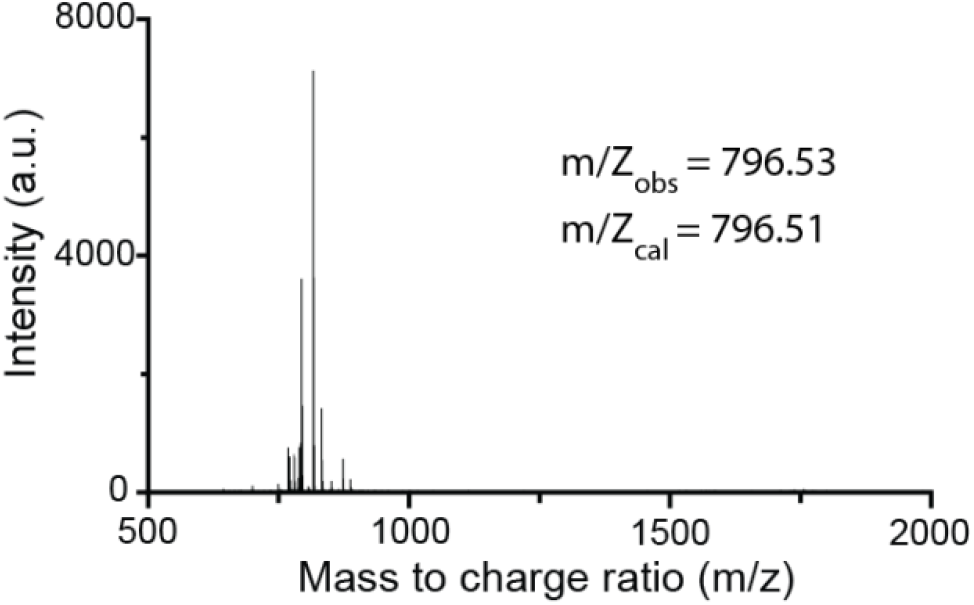
MALDI-ToF MS of synthesized IKVAV peptide.

**Figure S3:**
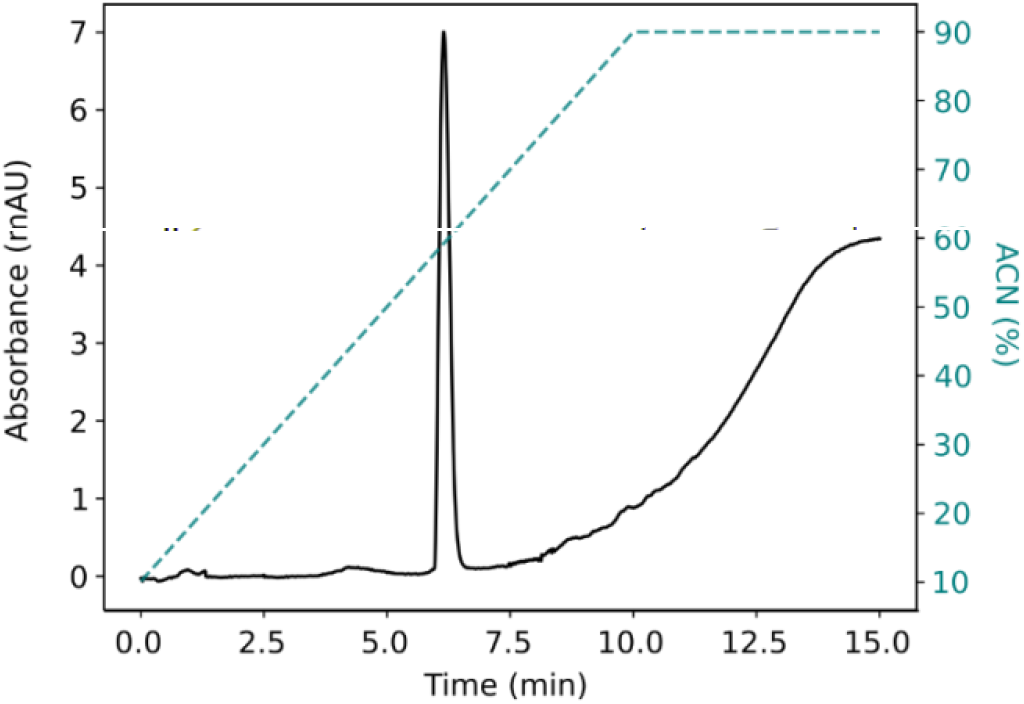
HPLC trace of synthesized IKVAV peptide ran on a C-18 column with 10-90% acetonitrile (can) gradient with UV absorbance measured at 229 nm

**Table S1a:**
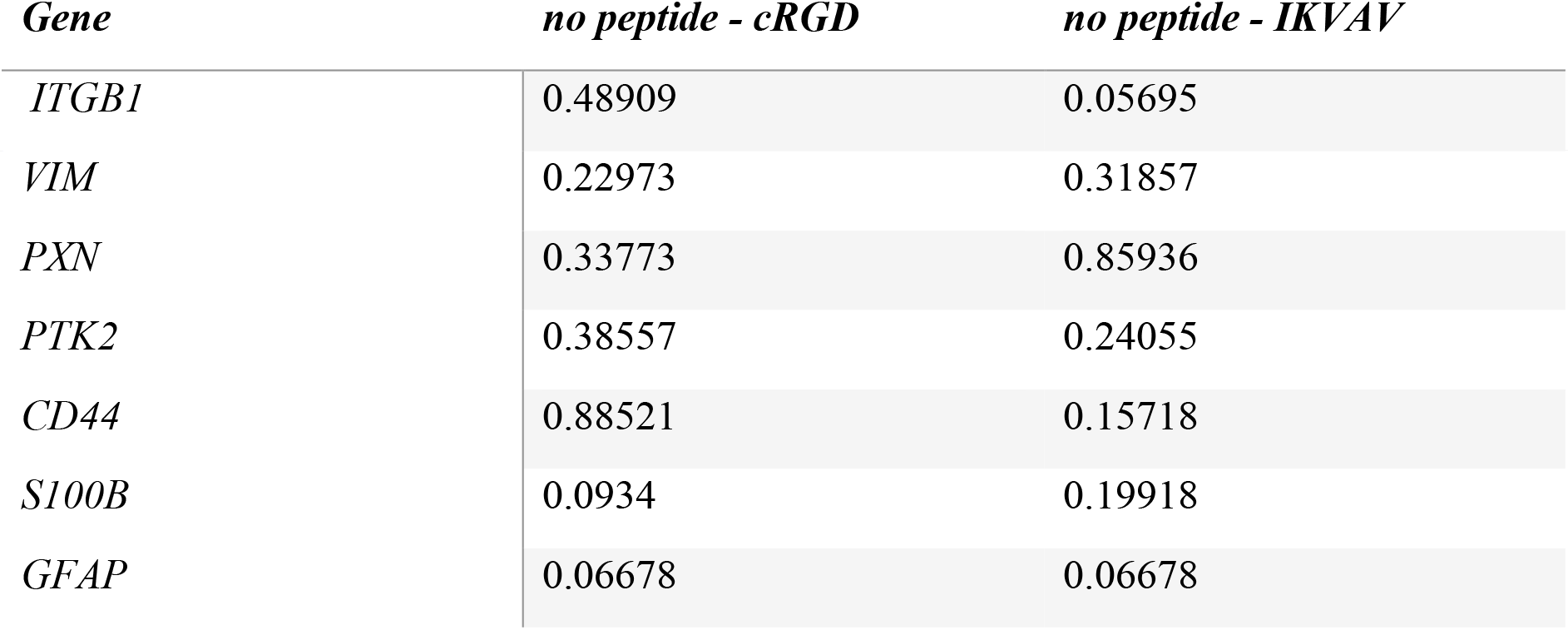
a) p-values for evaluating peptide effect on mRNA expression levels for FPA presented in Figure 4.

**Table S1b:**
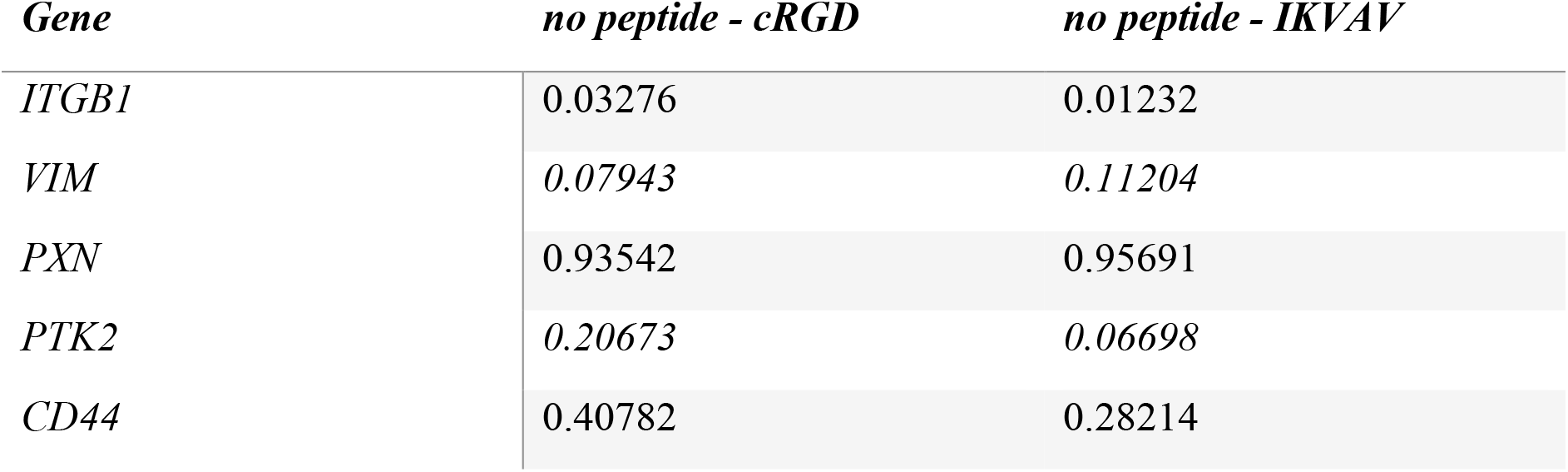
b) p-values for evaluating peptide effect on mRNA expression levels for U87, presented in Figure 5.

**Figure S4:**
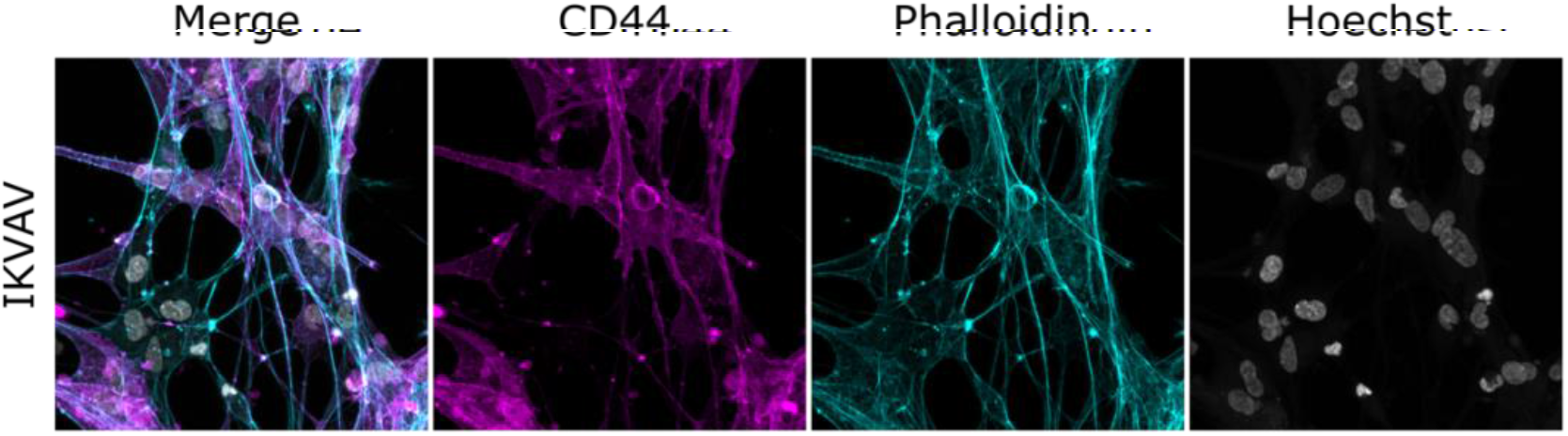
Confocal images of FPA encapsulated in IKVAV supplemented hydrogels.

